# Selective inhibition of MR1-restricted T cell activation by a novel MR1-targeting nanobody

**DOI:** 10.1101/2025.06.11.659204

**Authors:** Timothy Bates, Corinna Kulicke, Sintayehu Gurmessa, Se-Jin Kim, Sushant Suresh, Jules Reyes-Weinstein, Audrey Hinchliff, John Burke, David Lewinsohn, Fikadu G. Tafesse

## Abstract

MR1 is a non-polymorphic, ubiquitously expressed, MHC class I-like antigen-presenting molecule that presents small-molecule metabolites to T cells. Studies have shown that MR1 plays a role in microbial infection, inflammation, and tumor immunity. The antigens it presents include metabolites of microbial and self-origin as well as small-molecule drugs and form stable complexes with MR1 that are displayed on the cell surface to activate T cells. However, unlike classical MHC I and II molecules, the fundamental biology of MR1 remains poorly understood, particularly the mechanisms governing antigen loading and intracellular trafficking. This knowledge gap is largely due to the lack of molecular tools available to precisely manipulate MR1 function. In this study, we describe a high-affinity (1.6 nM *K*_*D*_) anti-MR1 nanobody, MR1Nb1. We characterize the binding of this nanobody including affinity by ELISA and kinetics by BLI. Crucially, we map the binding epitope of MR1Nb1 on MR1 by HDX-MS, providing key insights into the mechanism through which it blocks MR1T cell activation. In functional assays MR1Nb1 effectively and specifically blocks MAIT cell activation by cells infected with *M. tuberculosis* or treated with *M. smegmatis* supernatant. This nanobody represents a unique and versatile tool for the field, as it can be produced inexpensively and expressed intracellularly within antigen-presenting cells. Hence, our study provides a powerful new molecular probe for dissecting the mechanistic underpinnings of MR1 biology and uncover its broader roles in immunity.

## Introduction

Major histocompatibility complex (MHC) class I proteins are critical for CD8^+^ T cell activation in response to both microbial infections and cancer. MR1 is an MHC class I-like protein dedicated to sensing of small-molecule metabolites, including those derived from intermediates of the riboflavin biosynthesis pathway (1–3) and is widely expressed in human tissues (4). It is recognized by an “innate-like” subset of T cells known as mucosal-associated invariant T (MAIT) cells, which are abundant in epithelial tissues and broadly antimicrobial (5,6). Other MR1-restricted T cells (MR1T) have since been identified which recognize a broader range of antigens (7–9), of which MAIT cells appear to be an important and distinct subset. The activation of MAIT cells relies on engagement with MR1 on the surface of antigen presenting cells (APCs), and MR1 must be correctly loaded with ligand in order to reach the cell surface (10–12). Recent studies have demonstrated that MR1 antigen loading can occur through multiple mechanisms across various cellular compartments (Karamooz et al., 2023; Kulicke et al., 2024). However, the precise molecular details of antigen loading and subcellular trafficking remain incompletely understood.

The proper coordination of MAIT cells and MR1-bearing APCs is important for stimulation of appropriate immune responses to microbial pathogens as well as some vaccines (13). In particular, the role of MAIT cells has been extensively studied in the context of tuberculosis (TB) (14–16), the leading cause of death from a single infectious agent worldwide (17). A single nucleotide polymorphism (SNP) in MR1 which reduces expression has been associated with susceptibility to TB disease (18).

In addition to its well-established role in microbial infection, MR1 has also been implicated in cancer immunity (19,20). A recently characterized class of T cells, termed MR1T cells, can recognize antigens presented by MR1 even in the absence of infection (21). This recognition is primarily driven by self-antigens, specifically nucleobase adducts that bind to MR1 and stimulate MR1-restricted T cells. Carbonyl-modified nucleobase adducts have been detected within MR1 molecules on tumor cells, and their abundance and antigenicity are enhanced by drugs that promote carbonyl accumulation (22). More recently, the carbonyl nucleobase adduct M_3_ Ade has been identified as a potent antigen capable of activating adaptive polyclonal MR1-restricted T cells. Using MR1-M_3_ Ade tetramers, heterogeneous MR1-reactive T cell populations have been detected ex vivo in healthy individuals, patients with acute myeloid leukemia, and tumor-infiltrating lymphocytes from non-small cell lung adenocarcinoma and hepatocellular carcinoma (23). However, the molecular mechanisms that distinguish MR1 recognition of microbially derived versus self-derived metabolites remain incompletely understood. Developing tools to probe MR1 biology will be instrumental in advancing our understanding of its immunological functions in both infection and cancer.

Single-domain antibodies, also called nanobodies, are unique antibody fragments which can serve as highly versatile affinity reagents for a broad range of research activities. They are derived from heavy-chain-only antibodies which natively lack light chains, found in camelid species and cartilaginous fish (24,25). Because they are small (12-15 kDa) and composed of a single unglycosylated peptide chain, they can be efficiently produced in bacterial expression systems, and are readily engineered into fusion proteins (26,27). They are also able to fold and function in the cytoplasm of eukaryotic cells (28,29), enabling relatively simple generation of intracellular biosensors. Similarly, nanobodies can also be used to make inducible degron systems (30). Further, nanobody discovery can be performed using high throughput methods such as phage display, allowing for rapid generation of antigen-specific candidates (31–33).

In this study, we describe an alpaca-derived MR1-specific nanobody, MR1Nb1, which can block MAIT cell activation by *Mycobacterium tuberculosis* (Mtb)-infected APCs. We characterize the affinity and binding kinetics of this nanobody using enzyme-linked immunosorbent assay (ELISA) and bio-layer interferometry (BLI), perform epitope mapping with hydrogen-deuterium exchange (HDX), and test its functional activity in an enzyme-linked immunosorbent spot (ELISpot) assay with both *Mycobacterium Smegmatis* (*M. smegmatis*) supernatant and live Mtb infection. These tests each confirm that MR1Nb1 is an exceptionally high-affinity nanobody for human MR1 with strong inhibitory activity against MAIT cells. This tool will allow future studies to address new research questions about MR1 signaling and trafficking.

## Results

MR1 is widely expressed in human tissues, however its presentation and translocation to the cell surface requires association with β2-microglobulin (β2M) and loading with appropriate ligand (12) (Fig 1A). The complete repertoire of natural ligands loaded onto MR1 and recognized by MR1T cells during both health and disease are still a matter of active research, but several model ligands have been identified. These include 6-formylpterin (6-FP) which can be loaded into MR1 and bring it to the cell surface, but cannot activate MAIT cells; and 5-(2-oxopropylideneamino)-6-D-ribitylaminouracil (5-OP-RU) which is a highly potent ligand that both brings MR1 to the surface and activates MAIT cells (34,35). We utilized these two representative antigens in this study because of their widespread use and opposing effects, although other classes of MR1 antigens continue to be discovered (23).

**Figure 1.**
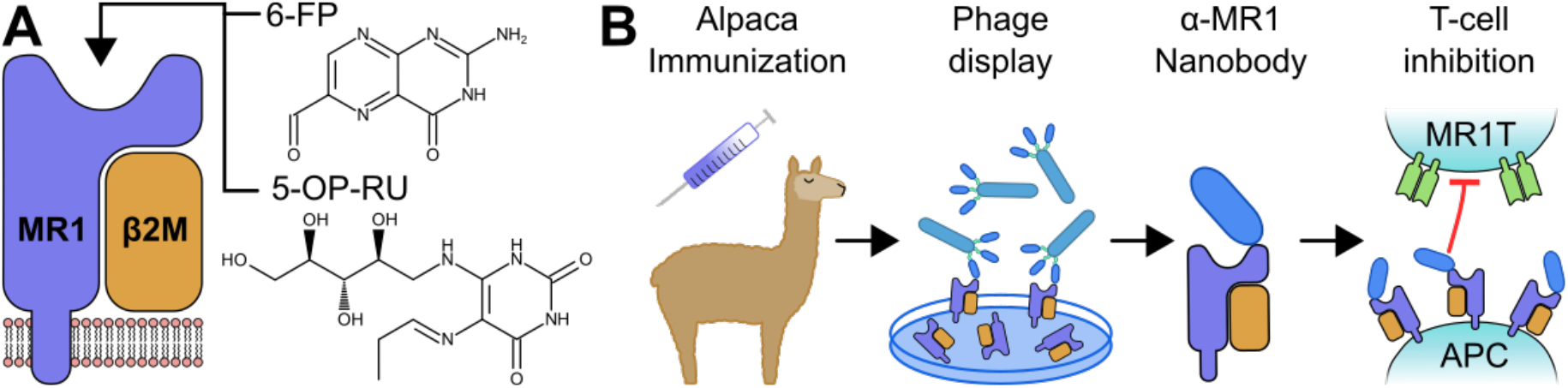
Schematic diagrams. Membrane-bound MR1 attached to its obligate binding partner, β2M. The antigen binding groove of MR1 can bind and display a variety of small molecule antigens including 6-FP and 5-OP-RU (A). Alpacas were immunized with recombinant MR1; phage display was performed on the resulting immune library; individual candidates were produced and characterized, including T-cell inhibition assays (B).

### Generation and characterization of MR1-specific nanobodies

We identified several MR1-specific nanobodies using phage display on an alpaca immune library (Fig 1B). We generated our library by immunizing an alpaca with recombinant MR1 protein. Following immunization, we collected blood and constructed a phage display library by cloning the VHH gene from isolated peripheral blood mononuclear cells (PBMCs). Then, to identify strong binders to MR1 in our phage library, we performed biopanning with MR1 and sequenced the resulting hits.

Upon expression and purification of these initial hits, we identified two nanobodies able to bind strongly to recombinant MR1, MR1Nb1 and MR1Nb3. We performed ELISA experiments to measure the 50% effective concentrations (EC_50_) for both of these nanobodies for recombinant MR1 protein loaded with either 6-FP or 5-OP-RU. These experiments showed an EC_50_ of 1.9 ng/mL and 4.8 ng/mL for MR1Nb1 on MR1 loaded with 6-FP and 5-OP-RU, respectively (Fig 2A,B). MR1Nb3 showed slightly higher EC_50_ values of 32 ng/mL and 69 ng/mL for MR1 loaded with 6-FP and 5-OP-RU, respectively.

**Figure 2.**
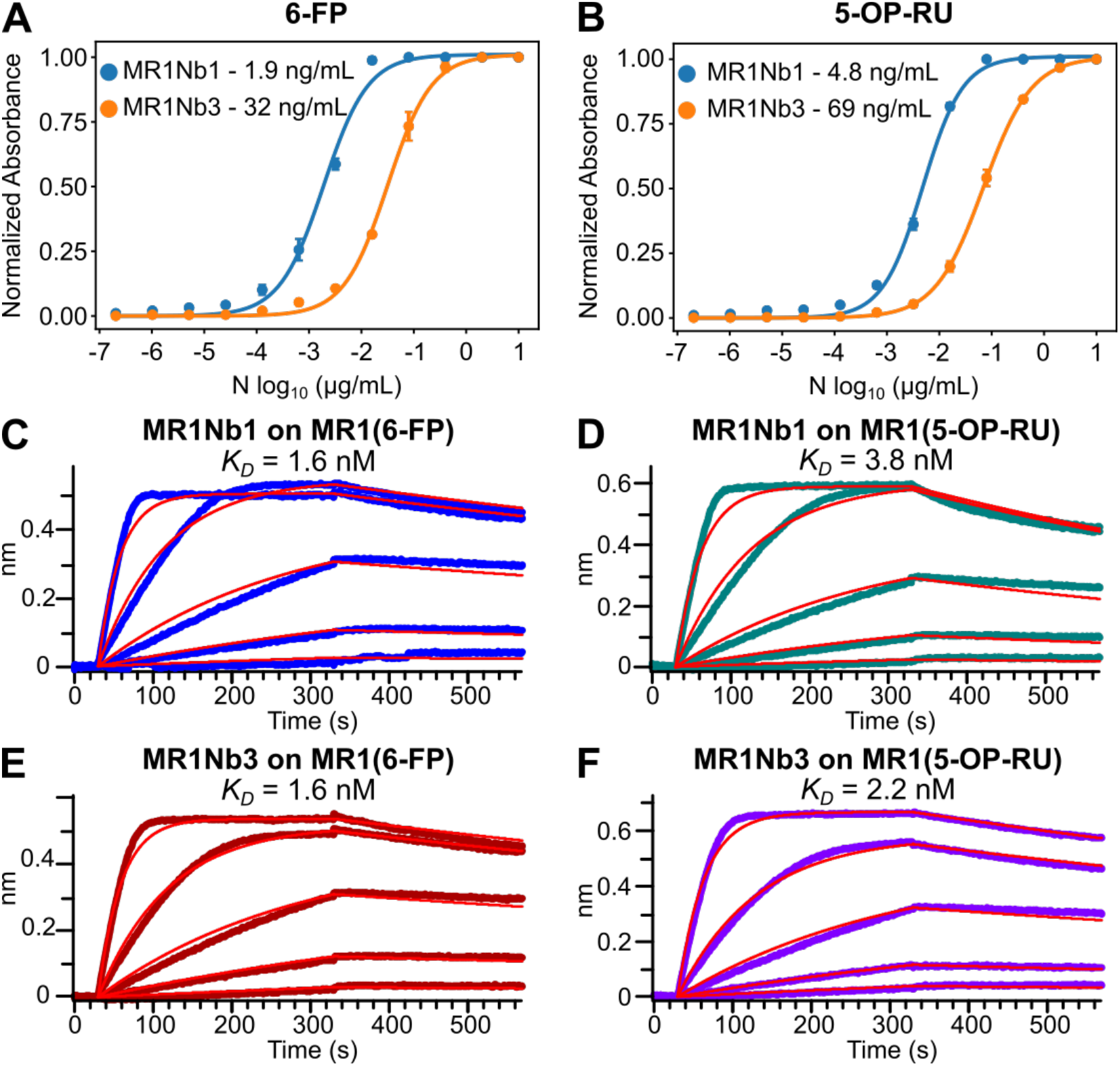
MR1Nb1 and MR1Nb3 bind to MR1 loaded with either 6-FP or 5-OP-RU. ELISA EC_50_ ;curves showing MR1Nb1 and MR1Nb3 binding to plates coated with 6-FP-loaded MR1 (A), and 5-OP-RU-loaded MR1 (B). Biolayer interferometry of MR1Nb1 on 6-FP-loaded MR1 (C), and 5-OP-RU-loaded MR1 (D); then similarly with MR1Nb3 on 6-FP-loaded MR1 (E), and 5-OP-RU-loaded MR1 (F). ELISA experiments were performed in triplicate (n=3) and BLI experiments were performed in duplicate (n=2).

Next, we wanted to establish the binding kinetics of MR1Nb1 and MR1Nb3. To do this, we performed BLI, which utilizes reflective biosensors to measure the rates of association and dissociation in real time (36). For these experiments, we used streptavidin-coated biosensors and loaded them with biotinylated MR1 loaded with either 6-FP or 5-OP-RU. MR1Nb1 displayed a *K*_*D*_ of 1.6 nM (*k*_*ON*_ 3.6×10^5^ M^-1^s^-1^; *k*_*OFF*_ 5.8×10^−4^ s^-1^) on 6-FP-loaded MR1 (Fig 2C), and a *K*_*D*_ of 3.8 nM (*k*_*ON*_ 3.0×10^5^ M^-1^s^-1^; *k*_*OFF*_; 1.1×10^−4^ s^-1^) on 5-OP-RU-loaded MR1 (Fig 2D). MR1Nb3 meanwhile, displayed a *K*_*D*_ of 1.6 nM (*k*_*ON*_ 3.4×10^5^ M^-1^s^-1^; *k*_*OFF*_ 5.4×10^−4^ s^-1^) on 6-FP-loaded MR1 (Fig 2E), and a *K*_*D*_ of 2.2 nM (*k*_*ON*_ 3.0×10^5^ M^-1^s^-1^; *k*_*OFF*_ 6.4×10^−4^ s^-1^) on 5-OP-RU-loaded MR1 (Fig 2F). For both VHHs, each concentration was run separately on fresh biosensors as regeneration was not possible due to an inability to completely wash off the bound nanobody with pH 1.7 glycine, pH 10 glycine, or 3M KCl.

### Epitope determination

As both VHHs showed excellent binding kinetics, we next sought to determine how they were binding. We again used BLI to set up a competition experiment to test whether MR1Nb1 and MR1Nb3 bind to distinct epitopes. We also compared MR1Nb1 and MR1Nb3 to the MR1-specific monoclonal antibody (mAb) 26.5 which is known to bind fully folded MR1 and inhibit MAIT cell activation (37).

We performed this competition experiment by first loading biosensors with MR1 and saturating with MR1Nb1 or MR1Nb3. We can then test for competition with the opposing nanobody or mAb 26.5 by observing whether they are still able to bind to the saturated MR1-nanobody complex. We found that MR1Nb3 could not bind to MR1Nb1-saturated MR1 while mAb 26.5 could still bind and similarly MR1Nb1 was unable to bind MR1Nb3-saturated MR1 while mAb 26.5 did bind (Fig 3A-D). This was true for both 6-FP- and 5-OP-RU-loaded MR1 and indicated that MR1Nb1 and MR1Nb3 likely bind overlapping epitopes, while mAb 26.5 binds at a distinct non-overlapping site on MR1. Because of their similarity in affinity and competitive binding, we decided to focus on MR1Nb1 in further experiments because of its better performance in our ELISA experiments.

**Figure 3.**
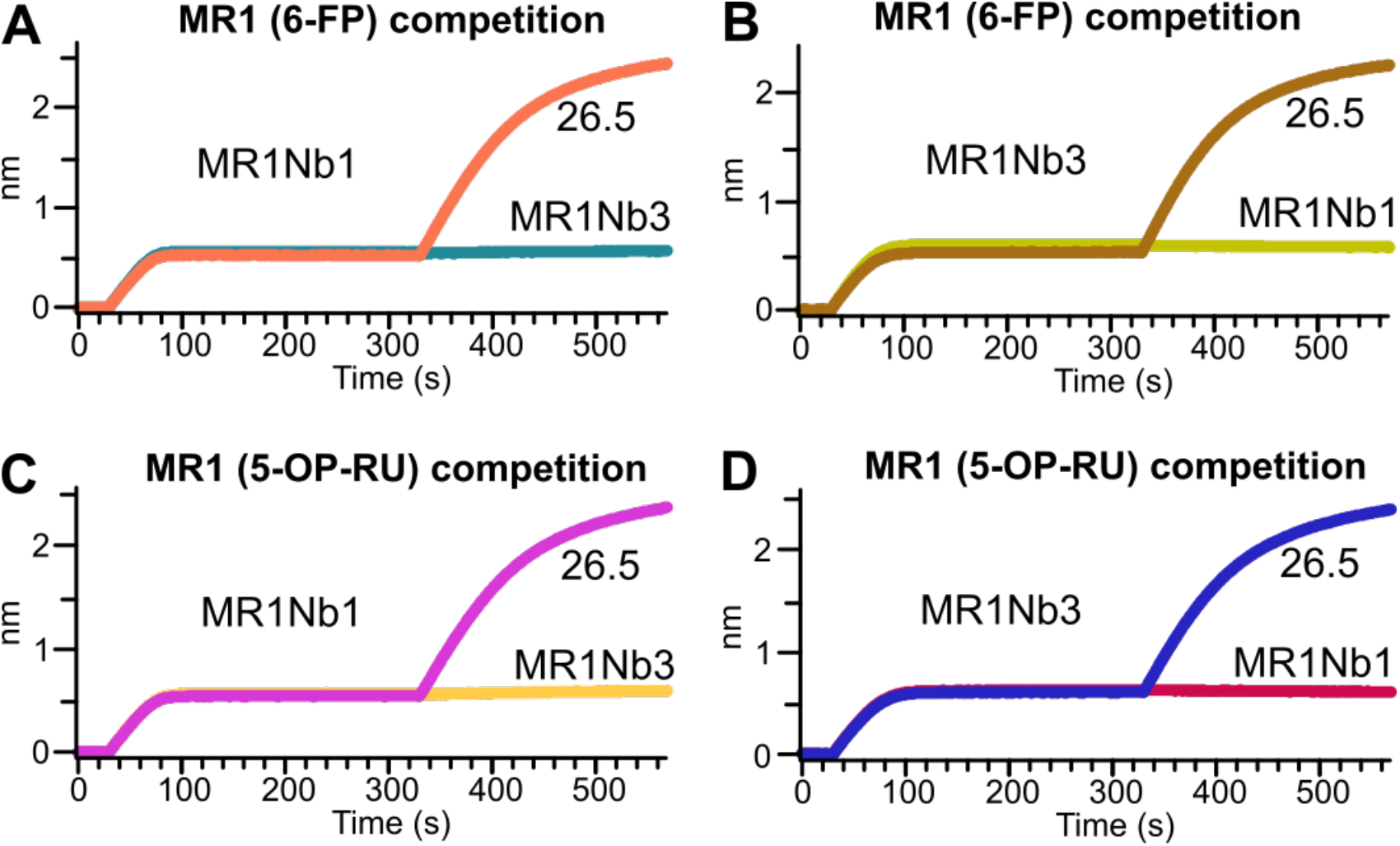
BLI showing competitive binding of MR1Nb1 with MR1Nb3, but not mAb 26.5. MR1 loaded with 6-FP were first bound with MR1Nb1 (A) or MR1Nb3 (B) then immediately transferred to the opposing VHH or mAb 26.5. This was repeated with MR1 loaded with 5-OP-RU for MR1Nb1 (C) and MR1Nb3 (D).

To precisely define the binding epitope of MR1Nb1 on the MR1 protein, we performed hydrogen-deuterium exchange mass spectrometry (HDX-MS). This technique maps the binding site of two proteins by observing the decrease in deuterium incorporation by residues with differential solvent exposure in bound and unbound states (38). Using this method, we compared apo and MR1Nb1-bound MR1 and found a significant decrease in deuterium incorporation between residues 141 and 165, indicating a likely site of interaction (Fig 4A). This epitope lies to one side of the ligand binding cleft of MR1, proximal to the reported binding site of some MAIT cell T cell receptors (TCRs) such as MAIT clone #6 from (39)(Fig 4B). In the areas with coverage, there were no other peptides with significant decreases in deuterium incorporation of more than 10% (Fig 4C), and the peptides which did show strong differences had consistent trends over time (Fig 4D).

**Figure 4.**
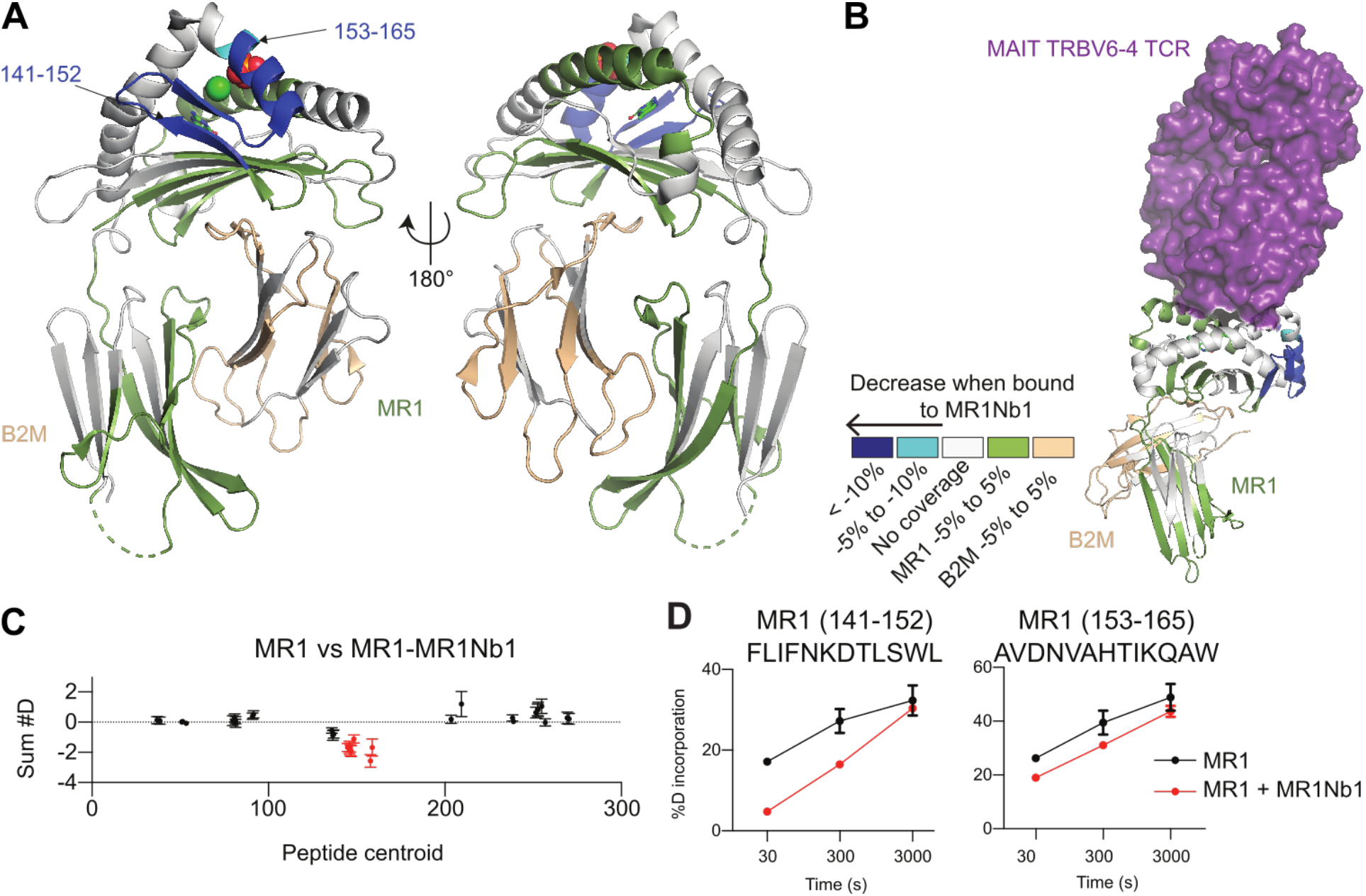
HDX-MS analysis of the interaction of MR1 with MR1Nb1. HDX differences (defined as >5%, 0.4 Da, and P< 0.01 in an unpaired two-tailed t test at any time point) upon MR1 binding to MR1Nb1 mapped on MR1 structure (PDB: 4GUP) (A). HDX differences of MR1 when bound to MR1Nb1 mapped on the structure of a TRBV6-4 MAIT TCR in complex with MR1 (PDB: 4PJ7) (B). Sum of the deuteron difference of MR1 upon binding to MR1Nb1 across the entire time course (C). Peptides that met the significance criteria described in (A) are colored red. Error is shown as the sum of SDs across all time points (n=3). Deuterium incorporation in MR1 peptides showing significant changes in the presence (red) and absence (black) of MR1Nb1 (D).

### Inhibition of MAIT cell activation by MR1Nb1

Next, we performed functional blocking studies to determine the ability of the generated nanobody to interfere with MR1-mediated antigen presentation. We used two T cell clones established in our laboratory, which are restricted by MR1 and the classical antigen presenting molecule human leukocyte antigen (HLA)-B45, respectively, and robustly produce interferon (IFN)-γ in response to APCs presenting exogenous antigens or infected with Mtb (2,40,41). This system allowed us to specifically assess the capacity of the nanobody to block MR1-dependent T cell activation without interfering with conventional HLA-mediated antigen presentation.

When using the airway epithelial cell line BEAS-2B as APCs, MR1Nb1 blocked MR1-mediated presentation of *M. smegmatis* supernatant, which is a source of exogenous MAIT cell antigens (10,42), with an efficiency similar to the routinely used anti-MR1 blocking antibody clone 26.5 (37,43) (Figure 5A). Importantly, neither MR1Nb1 nor antibody 26.5 impacted the presentation of peptide antigen to an HLA-B45-restricted classical T cell clone, confirming specificity for MR1 (Figure 5B). Similar results were obtained when using Mtb-infected BEAS-2B cells as the APCs (Figure 5C,D). While both MR1Nb1 and antibody 26.5 blocked the activation of the MR1-restricted MAIT cell clone, IFN-γ production by the classically restricted T cell clone was unaffected. Thus, the novel MR1 nanobody generated here is a valuable alternative to the currently available anti-MR1 antibodies for specifically and efficiently blocking MR1-mediated antigen presentation in functional experiments.

**Figure 5.**
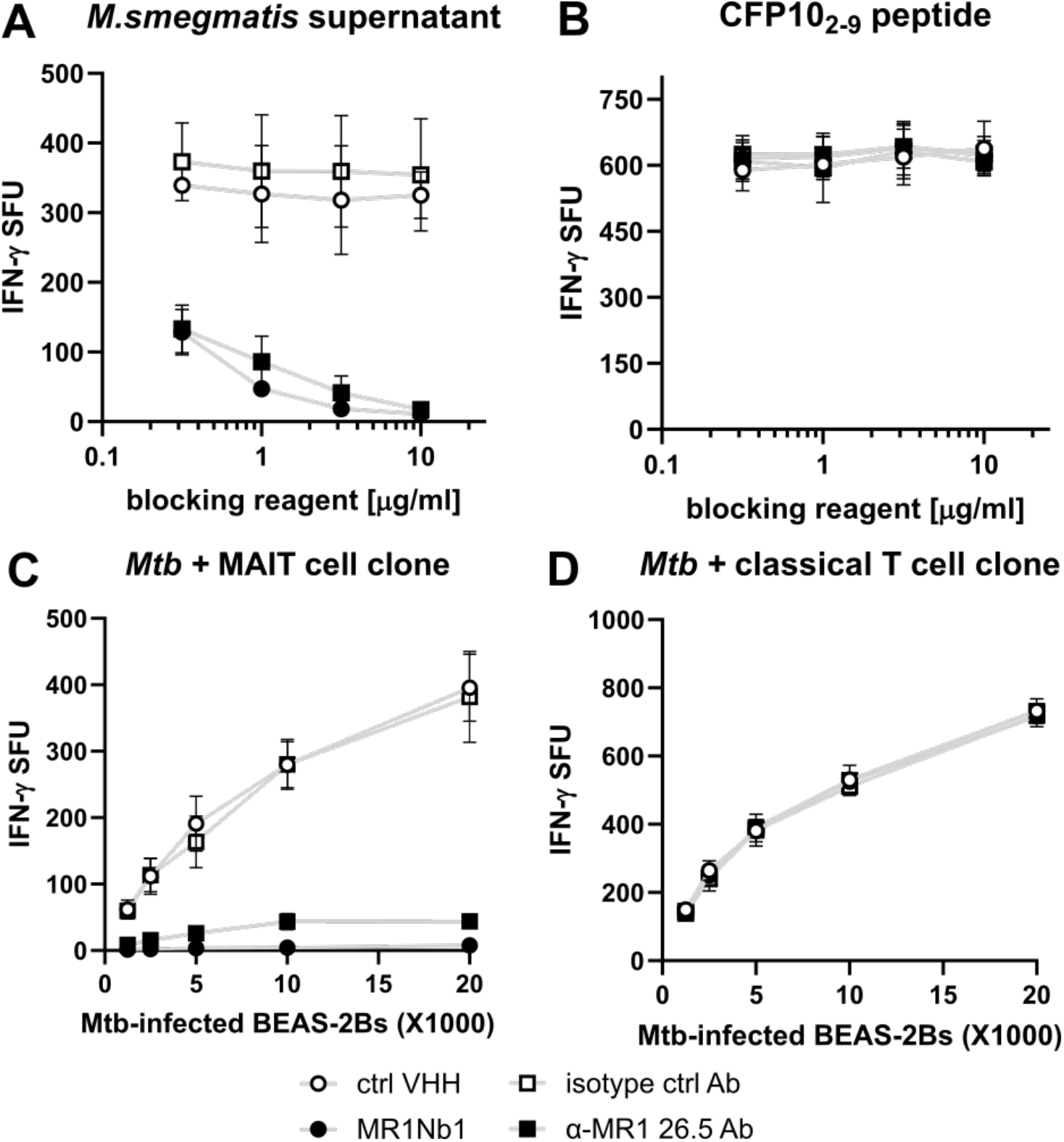
MR1Nb1 specifically blocks MR1-mediated presentation of both exogenous ligands and antigens derived from intracellular infection with Mtb. BEAS-2B cells were pre-treated with an irrelevant control nanobody (open circles), MR1Nb1 (filled circles), an IgG2a isotype control antibody (open squares), or the anti-MR1 antibody clone 26.5 (filled squares) at the indicated concentrations, incubated with either *M. smegmatis* supernatant (A) or CFP10_2-9_ peptide (B), and then co-cultured with the MR1-restricted MAIT cell clone D416-G11 (A) or the HLA-B45-restricted classical T cell clone D466-A10 (B) overnight. BEAS-2B cells were infected with Mtb strain H37Rv at an MOI of 8 overnight before incubation with 5 μg/ml of each blocking reagent and subsequent co-culture with MAIT cell clone D416-G11 (C) or classical T cell clone D466-A10 (D). IFN-γ production was measured as spot-forming units (SFU) by ELISpot. Data are the pooled means of technical duplicates from (n=3) independent experiments and shown as mean +/-standard deviation.

## Discussion

In this study, we present a nanobody with high affinity (*K*_*D*_ = 1.6 nM) to MR1 that blocks MAIT cell activation by Mtb-infected cells. As a monomer, its performance is similar to the existing bivalent mAb 26.5, though it appears to bind to a distinct non-overlapping epitope. It would be straightforward to produce a multivalent MR1Nb1 construct with improved binding affinity, if necessary for a particular application.

Like the TCR in the MR1-bound structure in Figure 4B (PDB: 4PJ7), the MAIT cell clone used in the ELISpot experiments in this study expresses the canonical MAIT TRAV1-2 TRAJ33 TCR alpha chain and the TRBV6-4 TCR beta chain (39,40). Thus, it presumably binds MR1 with a similar footprint, centrally above the antigen binding cleft. While the epitope for the anti-MR1 antibody 26.5 has not been unequivocally identified, several residues similarly located centrally on top of the ligand binding groove are likely important (37), indicating that antibody binding could sterically interfere with TCR binding.

In contrast, MR1Nb1 appears to bind MR1 adjacent to the antigen binding pocket and still blocks engagement by the MAIT cell TCR. While we have positively identified regions of engagement, it is possible that additional contact sites are present in the areas without coverage in our HDX-MS data. Based on the available data, it seems likely that a mechanism other than direct steric interference is responsible for blocking TCR engagement, but residue level information about these interactions would require a higher resolution mapping of the epitope using X-ray crystallography or Cryo-EM. Regardless, MR1Nb1 displays high-affinity binding to MR1 which is stable under acidic, basic, and high-salt conditions.

The role of MR1 in microbial pathogenesis is well established, particularly in presenting vitamin B metabolite-derived antigens to MAIT cells during infection. More recently, growing evidence indicates that MR1 also plays a significant role in cancer immunity (19,20). Notably, the carbonyl nucleobase adduct M3Ade has been identified as a self-antigen capable of activating adaptive, polyclonal MR1T cells. Correspondingly, MR1-reactive T cell populations have been detected in healthy individuals, patients with acute myeloid leukemia, and within tumor-infiltrating lymphocytes from lung adenocarcinoma and hepatocellular carcinoma (23). Given these diverse roles of MR1 in infection and tumor immunity, it is likely that our nanobody will modulate MR1-dependent T cell activation not only during microbial infection but also within the tumor microenvironment. This opens the possibility of using MR1-targeting nanobodies as tools to investigate and potentially manipulate MR1-restricted immunity in cancer.

We envision that MR1Nb1 will serve as a versatile tool for dissecting MR1 biology, particularly its intracellular trafficking and antigen presentation dynamics. Its ease of production in *E. coli* and modular nature make it highly amenable to protein engineering. MR1Nb1 can be reformatted into multivalent constructs to enhance avidity or modified into bispecific formats that simultaneously engage MR1 and other surface markers, enabling targeted modulation of antigen-presenting cells. Moreover, its small size and stability make it ideal for expression within specific subcellular compartments, allowing researchers to track MR1 trafficking in live cells with spatial precision or to intrinsically modulate MAIT cell activation without disrupting endogenous MR1 expression. In future studies, nanobodies with complementary properties could be developed, such as those that bind MR1 without blocking T cell activation, or chaperone-like nanobodies that stabilize MR1 and promote ligand-independent surface expression. Together, these nanobody-based tools offer exciting opportunities to probe MR1 function and manipulate MR1-restricted immunity in a highly specific and tunable manner.

## Methods

### Ethics Declaration

This study was conducted according to the principles expressed in the Declaration of Helsinki. Study participants, protocols, and consent forms were approved by the Institutional Review Board at Oregon Health & Science University (OHSU) (IRB00000186). All ethical regulations relevant to human research participants were followed. Peripheral blood mononuclear cells (PBMC) and human serum from human subjects were only used to expand T cell clones and in ELISpot medium as described in the methods section. All findings in this manuscript are based on T cell clone function or on other cell line function, e.g. BEAS-2B cell line. No data is presented on the PBMC from human subject participants and sex and gender were not taken into consideration.

### Phage display

Phage display was performed as previously described (33). Briefly, Alpacas were immunized 3 times, 3 weeks apart with recombinant MR1 protein provided as a generous gift by Dr. Erin Adams (UChicago), then whole blood was collected 1 week following the final immunization. PBMCs were isolated using Ficoll-Paque PLUS and preserved with RNAlater. RNA was extracted with a Qiagen RNeasy kit and cDNA was produced using Superscript III. PCR was performed to amplify VHH genes and cloned into a phagemid vector and transformed into TG1 *E. coli*. The bacterial library was then infected with M13 helper phage to produce the phage library, which was purified by PEG precipitation and resuspended in PBS. Panning plates were prepared by coating with 2 μg/mL recombinant MR1 protein, washed with PBST (0.1% Tween-20), blocked with 2% BSA in PBST and bound with the purified phage library in blocking buffer for 1 hour at room temperature (RT). The plates were then washed 10 times with PBST, eluted with 200 mM glycine pH 2.2, and neutralized with 1 M Tris pH 9.1 before infecting log-phase ER2738 *E. coli* and plating on agar plates. Individual colonies were picked from these plates and sent for sequencing.

### VHH purification

Selected VHHs were expressed and purified as previously described (33). Briefly, VHH genes were cloned into a pHEN vector with a C-terminal LPETG sortase tag followed by a 6x His tag. VHH plasmids were transformed into WK6 *E. coli* and grown in Terrific Broth and induced with IPTG overnight at 30°C. VHHs were purified from the periplasm by osmotic shock and bound to Ni-NTA beads before washing with 50 mM HEPES, 150 mM NaCl, pH 8.0, and eluting with wash buffer plus 500 mM Imidazole. Purified VHHs were then dialyzed into PBS and stored at -80°C.

### ELISA

Purified recombinant MR1 protein loaded with 6-FP or 5-OP-RU was obtained from the NIH tetramer core (1).

ELISA plates (Maxisorp) were coated with 2 μg/mL purified recombinant MR1 protein overnight at 4°C. Plates were blocked in wash buffer (1% BSA, 1% PVP, 0.1% Tween-20 in PBS) for 30 minutes at RT. Five-fold dilutions starting from 10 μg/mL of VHH were incubated for 1 hour at RT. Plates were washed with PBST three times between each antibody addition. Biotinylated-anti-VHH antibody and streptavidin-HRP secondary antibodies were used at a dilution of 1:10,000 in blocking buffer for 1 hour each at RT. Plates were developed with OPD substrate and read at 492 nm using a CLARIOstar Plus plate reader (BMG).

### BLI

BLI experiments were performed using previously described methods (36), using purified recombinant MR1 protein loaded with 6-FP or 5-OP-RU was obtained from the NIH tetramer core (1). Briefly, streptavidin (SA) biosensors were soaked in deionized water for 30 minutes prior to use. Solutions of antigen and nanobody were prepared in running buffer (RB: 10 mM HEPES, 150 mM NaCl, 3mM EDTA, 0.005% Tween-20, 0.1% BSA, pH 7.5). For each experiment, pre-soaked biosensors were blocked with RB, loaded with MR1, blank baseline in RB, blank association in RB, blank dissociation in RB, baseline in RB, association with nanobody or mAb, dissociation in RB. Nanobodies were tested at 0, 1, 3.16, 10, 31.6, and 100 nM, each on separate biosensors. Data were double-reference subtracted for kinetics calculations using Data Analysis HT (v10.0, Sartorius) using a 1:1 binding model and global fitting.

### HDX-MS sample preparation

HDX reactions used purified recombinant 6-FP-loaded MR1 protein from the NIH tetramer core (1). examining MR1 in the presence and absence of MR1Nb1 were carried out in 20 μL reaction volumes containing 20 pmol of MR1 and 40 pmol of MR1Nb1 (1 μM MR1, 2 μM MR1Nb1 in the bound state). The exchange reactions were initiated by the addition of 16 μL of D_2_;O buffer (50 mM HEPES pH 7.5, 150 mM NaCl) to 4 μL of protein (final D_2_O concentration of 74% [v/v]). Reactions proceeded for 3s, 30s, 300s, and 3000s at 18°C before being quenched with ice cold acidic quench buffer (0.7M TCEP, 2M Guanidine HCl, 2% formic acid) resulting in a final concentration of 0.25 M TCEP, 0.7 M guanidine HCl and 0.7% formic acid at pH 2.5. All conditions and timepoints were created and run in independent triplicate. All samples were flash frozen immediately after quenching and stored at -80°C.

### Protein digestion and MS/MS data collection

Protein samples were rapidly thawed and injected onto an integrated fluidics system containing a HDx-3 PAL liquid handling robot and climate-controlled (2°C) chromatography system (Trajan), an ACQUITY UPLC I-Class Series System (Waters), as well as an Impact HD QTOF Mass spectrometer (Bruker). The full details of the automated LC system have been previously described (44). Samples were run over an immobilized pepsin column (Affipro; AP-PC-001) at 200 μL/min for 4 minutes at 2°C. The resulting peptides were collected and desalted on a C18 trap column (ACQUITY UPLC BEH C18 1.7 μm column, 2.1 mm x 5 mm; Waters 186004629). The trap was subsequently eluted in line with an ACQUITY 300Å 1.7 μm particle, 100 mm × 2.1 mm BEH C18 UPLC column (Waters; 186003686), using a gradient of 3-10% B (Buffer A 0.1% formic acid; Buffer B 100% acetonitrile) over 1.5 minutes, followed by a gradient of 10-25% B over 4.5 minutes, followed by a gradient of 25-35% B over 5 minutes, finally after 1 minute at 35% B a gradient of 35-80% B over 1 minute was used. Mass spectrometry experiments were acquired over a mass range from 150 to 2200 m/z using an electrospray ionization source operated at a temperature of 200°C and a spray voltage of 4.5 kV.

### Peptide identification

Peptides were identified from non-deuterated samples of MR1 using data-dependent acquisition following tandem MS/MS experiments (0.5s precursor scan from 150-2000 m/z; twelve 0.25s fragment scans from 150-2000 m/z). The MS/MS datasets were analysed using FragPipe v18.0 and peptide identification was carried out by using a false discovery-based approach using a database of purified proteins and known contaminants (45–47). MSFragger was used, and the precursor mass tolerance error was set to -20 to 20 ppm. The fragment mass tolerance was set at 20 ppm. Protein digestion was set as nonspecific, searching between lengths of 4 and 50 aa, with a mass range of 400 to 5000 Da.

### Mass Analysis of Peptide Centroids and Measurement of Deuterium Incorporation

HD-Examiner Software (Sierra Analytics) was used to automatically calculate the level of deuterium incorporation into each peptide. All peptides were manually inspected for correct charge state, correct retention time, appropriate selection of isotopic distribution, etc. Deuteration levels were calculated using the centroid of the experimental isotope clusters. Results are presented as relative levels of deuterium incorporation, and the only control for back exchange was the level of deuterium present in the buffer (74%). Differences in exchange in a peptide were considered significant if they met all three of the following criteria: ≥5% change in exchange, ≥0.45 Da difference in exchange, and a p value <0.01 using a two tailed student t-test. The entire HDX-MS dataset with all the values and statistics are provided in the source data. Samples were only compared within a single experiment and were never compared to experiments completed at a different time with a different final D_2_O level. The data analysis statistics for all HDX-MS experiments are provided in the source data according to the guidelines of (38). HDX-MS proteomics data generated in this study have been deposited to the ProteomeXchange Consortium via the PRIDE partner repository with the dataset identifier PXD058493 (48).

### Cells

BEAS-2B cells were obtained from ATCC and cultured in Dulbecco’s Modified Eagle Medium (gibco or Corning) supplemented with 10% heat-inactivated fetal bovine serum (GeminiBio) and 2% L-Glutamine (gibco). T cell clones D426-G11 and D466-A10 have been described previously and were expanded and maintained as before (2,43). D426-G11 is a TRAV1-2^+^ MAIT cell clone derived from a latently Mtb-infected individual (2,40,43). D466-A10 was derived from an individual with active tuberculosis and recognizes CFP10 peptide 2-9 in the context of HLA-B45 (41). *M. smegmatis* supernatant as a source of exogenous MAIT cell antigens was generated by growing *M. smegmatis* (strain mc^2^155) in Middlebrook 7H9 broth supplemented with Middlebrook ADC Enrichment (both BD), 0.5% Glycerol, and 0.05% Tween-80 (EMD Chemicals) at 37°C with shaking, followed by centrifugation and filtration of the supernatant through a 0.22 μm filter (Millipore). Aliquots were frozen at -80°C and thawed immediately before using. Mtb H37Rv obtained from ATCC was grown in Middlebrook 7H9 Broth supplemented with Middlebrook ADC (BD), 0.05% Tween-80 (OmniPur), and 0.5% glycerol (Thermo Fisher Scientific). Titer plates were used to determine the titer of an Mtb stock and frozen aliquots were stored at -80°C. After thawing, Mtb was passaged 20 times through a tuberculin syringe (BD) before infection.

### ELISpot assays

96-well mixed cellulose esters enzyme-linked immunosorbent spot (ELISpot) plates (Millipore) were coated with IFN-γ capture antibody clone 1-D1K (Mabtech) in 0.1 M Na2CO3, 0.1 M NaHCO3, pH=9.6 at 4°C overnight. Plates were washed three times with sterile phosphate buffered saline (PBS; gibco or Corning) and blocked with Roswell Park Memorial Institute (RPMI) 1640 (gibco or Corning) supplemented with 10% heat-inactivated human serum, 2% L-Glutamine, and 0.1% gentamycin (gibco) for at least one hour at room temperature. Blocked plates were either used immediately or stored at 4°C. For ELISpots with exogenous ligands, 1e^4^ BEAS-2B cells were plated in each well of the ELISpot plate and incubated with the indicated concentrations of control VHH [saRBD-1 (33)], MR1 MR1Nb1, IgG2a control antibody clone MOPC-173 (BioLegend), or purified anti-MR1 antibody 26.5 (OHSU antibody core) for at least one hour at 37°C. VHHs were sterile filtered through 0.22 μm centrifugal filter units (Millipore) and quantified by BCA assay (Thermo Scientific) before use in ELISpots. 2.5 μl/well of *M. smegmatis* supernatant or 20 μg/well of CFP10_2-9_ peptide (GeneMed) were then added, and plates were incubated for at least another hour at 37°C before addition of 2.5×10^3^ T cells/well. MR1-restricted D426-G11 T cells were added to the *M. smegmatis* supernatant wells and HLA-B45-restricted D466-A10 T cells were added to the peptide wells. Cells were co-cultured at 37°C for at least 18h. Plates were washed extensively in PBS (Sigma Aldrich) + 0.05% Tween-20 (Affymetrix) before incubation with alkaline phosphatase-conjugated secondary antibody clone 7-B6-1-ALP (Mabtech) in PBS supplemented with 0.5% BSA (Fisher Scientific) and 0.05% Tween-20 for two hours at room temperature. Plates were washed in PBS + 0.05% Tween-20 again and developed using BCIP/NBT-plus developer (Mabtech). IFN-γ spots were enumerated using an AID ELISpot reader and AID ELISpot Software Version 7 (Autoimmun Diagnostika GMBH).

For ELISpots using Mtb-infected cells, 4×10^5^ BEAS-2B cells/well were plated in a 6-well plate and left to adhere for at least five hours. Cells were then infected with freshly thawed Mtb H37Rv at an MOI of 8 overnight. Infected BEAS-2B cells were resuspended to 2×10^5^, plated in duplicate at 100 μl/well for each serial dilution and incubated with each blocking reagent at a final concentration of 5 μg/ml for one hour at 37°C. 1×10^4^ of MR1-restricted D426-G11 T cells or HLA-B45-restricted D466-A10 T cells were added in 50 μl/well and co-cultured with Mtb-infected BEAS-2B cells at 37°C for at least 18h. ELISpot plates were washed, developed, and enumerated as described above.

## Data Analysis

ELISA data were analyzed and plotted using a previously published python script (49). ELISpot data were plotted and analyzed using GraphPad Prism Version 8.4.3.

## Acknowledgements

We would like to thank the participants who gave time and dedication to this health research as well as Erin Merrifield, Department of Pediatrics, OHSU, for her contributions to this study. The MR1 tetramer technology was developed jointly by Dr. James McCluskey, Dr. Jamie Rossjohn, and Dr. David Fairlie, and the material was produced by the NIH Tetramer Core Facility as permitted to be distributed by the University of Melbourne. Additional MR1 protein was a generous gift from Dr. Erin Adams. BLI data were generated on an Octet Red 384, which is made available and supported by the Oregon Health & Science University Biophysics Shared Resources Core and equipment grant number S10OD023413. We acknowledge the assistance of the Oregon Clinical & Translational Research Institute, which is supported by the National Center for Advancing Translational Sciences, National Institutes of Health, through Grant Award Number UL1TR002369.

The contents do not represent the views of the U.S. Department of Veterans Affairs or the United States Government.

## Author contributions

Conceptualization; T.A.B., C.A.K., D.M.L., F.G.T. Methodology; T.A.B., C.A.K., S.G., S.J.K., S.S., J.B.W., A.H. Formal analysis; T.A.B., C.A.K., S.S. Investigation; T.A.B., C.A.K., S.G., S.J.K., S.S., J.B.W., A.H. Resources; J.E.B., D.M.L., F.G.T. Writing – Original Draft; T.A.B. Writing – Review & Editing; All authors. Visualization; T.A.B., C.A.K., S.S. Supervision; T.A.B., J.E.B., D.M.L., F.G.T. Funding acquisition J.E.B., D.M.L., F.G.T.

